# Gaze when walking to grasp an object in the presence of obstacles

**DOI:** 10.1101/2024.11.20.624440

**Authors:** Dimitris Voudouris, Eli Brenner

## Abstract

People generally look at positions that are important for their current actions, such as objects they intend to grasp. What if there are obstacles on their path to such objects? We asked participants to walk into a room and pour the contents of a cup placed on a table into another cup elsewhere on the table. There were two small obstacles on the floor between the door and the table. There was a third obstacle on the table near the target cup. Participants mainly looked at the items on the table, but as they approached and entered the room they often looked at the floor near the obstacles, although there was nothing particularly informative to see there. They relied on peripheral vision and memory of where they had seen obstacles to avoid kicking the obstacles. From well before participants crossed the obstacles, they primarily looked at the object that they intended to grasp. We conclude that people look at positions at which they plan to interact with the environment in a specific manner, rather than at items that constrain such interactions.

## Introduction

When performing their daily actions, humans typically direct their gaze to positions of interest for the ongoing or upcoming movement. This is well documented for various tasks, from isolated pointing (Neggers & Bekkering, 2000) and grasping (Brouwer et al., 2009) movements, to the sequential actions that are required to prepare breakfast (Land et al., 1999), take part in sports (Mann et al., 2013), steer a car (Tuhkanen et al., 2021) and navigate staircases (Ghiani et al., 2023) or rough terrain (Matthis et al., 2018). Gaze shifts bring relevant positions of the environment onto the fovea, which is the part of the retina with the highest spatial resolution. People typically direct their gaze at the target of their movement before initiating a movement toward that target (Johansson et al., 2001; Hollands & Marple-Horvat, 2001), and keep fixating the target until reaching it (Brenner & Smeets, 2007; Cámara et al., 2018), which allows them to use the best possible visual information to both plan and guide the movement to the target. Relying on visual input with the highest possible spatial resolution is presumably important in demanding tasks that require high spatial and temporal precision, such as grasping an object (Voudouris et al., 2016) or intercepting moving objects (Brenner & Smeets, 2011).

In many tasks it is evident where the relevant information is to be found, but there are also tasks in which there are multiple important positions. An example is when simultaneously reaching with both hands to different targets, in which case people look at the target that requires a higher degree of visual guidance, such as the smaller target or the target that one is reaching for with the non-dominant hand (Riek et al., 2003). When reaching to grasp a single object in the presence of obstacles, humans primarily fixate the target object rather than the obstacle (Grant, 2015; Marotta & Graham, 2016; Voudouris et al., 2016), suggesting that it is important to have precise information about the target object, while obstacles can be cleared without looking directly at them, possibly because peripheral vision is precise enough to monitor obstacles’ positions relative to the moving arm. When tracking or intercepting targets with a cursor, people are known to look at the target rather than the cursor (Cámara et al., 2018; Danion & Flanagan, 2018), relying on peripheral visual information and kinaesthetic information to guide the cursor to the target. That humans can perform tasks without looking at all relevant positions is evident from the fact that they can successfully climb staircases while looking at their phones rather than at the steps, though they do slow down their stair climbing to do so (Ioannidou et al., 2017). When navigating flat terrains, humans often look straight ahead, away from potential footholds, but on rough, more complex terrain, gaze is primarily directed to upcoming footholds (Matthis et al., 2018; ‘t Hart et al., 2013). Similarly, when people need to step over obstacles of different heights while walking, they spend more time looking at the obstacles when the obstacles are higher (Patla & Vikers, 1997), showing that as the stepping accuracy demands increase, people rely more on central vision to guide their leg. All in all, there appears to be a close coupling between gaze allocation and body movements during continuous actions, whereby gaze is directed in the manner that best ensures that all aspects of the action are performed successfully.

We here examine gaze patterns when people walk into a room to grasp an object. In particular, we examine how gaze is influenced by obstacles on their walking path. We also examine how gaze is influenced by obstacles near the target object. Does making it more difficult to reach out and grasp the target object without colliding with an obstacle beside it make people keep their gaze close to those two items? If so, they will spend more time looking at the target object and surrounding obstacle, and therefore less time looking at the floor. They will therefore have to rely more heavily on memory of the obstacles’ positions and on peripheral vision to guide their steps to upcoming footholds and away from the obstacles along the walking path. We wanted to know whether people primarily fixate the target object, relying on peripheral vision to guide their foot placement, or whether they also often fixate obstacles along their walking path or the places at which they intend to place their feet. Previous work has shown that humans sequentially look at objects that they need to reach and at obstacles that they need to avoid when walking through a virtual scene with many objects and obstacles along their path (Tong et al., 2017). Thus, we might expect participants to fixate each obstacle as they approach it, only fixating the object that they need to grasp after having passed the floor obstacles, especially since the object that they need to grasp is always at the same position in the room.

Participants walked into a room to grasp a cup of water placed at a fixed position on a table. There were two obstacles (other cups) on the floor along the walking path, and a third obstacle (yet another cup) on the table near the target object. Upon grasping the cup of water, participants had to empty its contents into a final cup that could be at either end of the table. If gaze is allocated sequentially, participants should look at positions of interest as these positions become relevant. Thus, participants should first look at the first and then at the second obstacle on the floor to plan how to best avoid them. After crossing these obstacles, they should look at the obstacle next to the cup of water and the cup of water itself to plan the most suitable grasping posture. Finally, they should look at the cup into which they are going to pour the water. However, there are several alternative credible gaze patterns. One is that participants look at all relevant positions as soon as they can, for instance by quickly looking at both the obstacles on the floor as well as at the one on the table to determine the most efficient walking path and grasping posture considering all constraints imposed by the obstacles. Another is that they quickly direct and maintain their gaze at the cup of water and the surrounding obstacle, the interaction with which requires the highest precision, and rely on peripheral vision to avoid kicking over the obstacles on the floor. A sequential pattern has been found in continuous tasks such as walking (Tong et al., 2017), while maintaining gaze where most precision is required is the typical behavior in discrete tasks such as grasping (Voudouris et al., 2018). Most studies examining gaze allocation during grasping had participants sit or stand at a position from which they only had to move their arm to reach and grasp the object (Brouwer et al., 2009; Grant, 2019; Voudouris et al., 2016; Voudouris et al., 2018). Here, we examine how gaze is allocated when people have to take a few steps to grasp an object. Will they still mainly look at the object they intend to grasp, or will they first look at the obstacles on the floor?

## Methods

### Participants

Ten participants (6 women, age range: 21-33 median age: 26) volunteered to take part in the experiment. They had normal or corrected-to-normal vision, were right-handed by self-report, and free from any known neurological or musculoskeletal issues that would make it difficult for them to walk, or to grasp and manipulate a cup. Participants signed informed consent forms prior to taking part in the experiment. The study was approved by the local ethics committee of Justus Liebig University Giessen and was conducted in accordance with the Declaration of Helsinki (2013; except for §35, pre-registration). Participants were compensated with 8€/hour or with course credits for their efforts.

### Setup and data collection

Participants were asked to walk into a room (296 × 480 cm), pick up a cup of water placed on a table (161 × 80 cm), and pour the water into another cup placed at one of two corners of the table. They were instructed to do so without knocking over three obstacles, two of which were placed on the floor along their walking path and a third one near the target cup. The target cup (the cup with the water) was placed centrally on the table, 22 cm from the edge that was closest to the participant. The destination cup was placed 66.5 cm to the left or right of the target cup and at 11 cm from the edge of the table that was closest to the participant. Participants had to grasp and manipulate the cup using a precision grip with their right hand. The table was positioned so that its near edge was 280 cm from the entrance to the room. The two floor obstacles were placed inside the room and were positioned at 140 and 205 cm from the door. Each floor obstacle could be placed at one of two lateral positions that were 55 cm apart. The table obstacle was placed 11 cm to the side of and 11 cm in front of the target cup (aligned with the destination cup). All objects were plastic cups (15 cm tall, 9 cm mouth diameter, 5.5 cm base diameter) of different colors. A schematic depiction of the setup and of the possible object configurations is shown in Figure 1.

**Figure 1.**
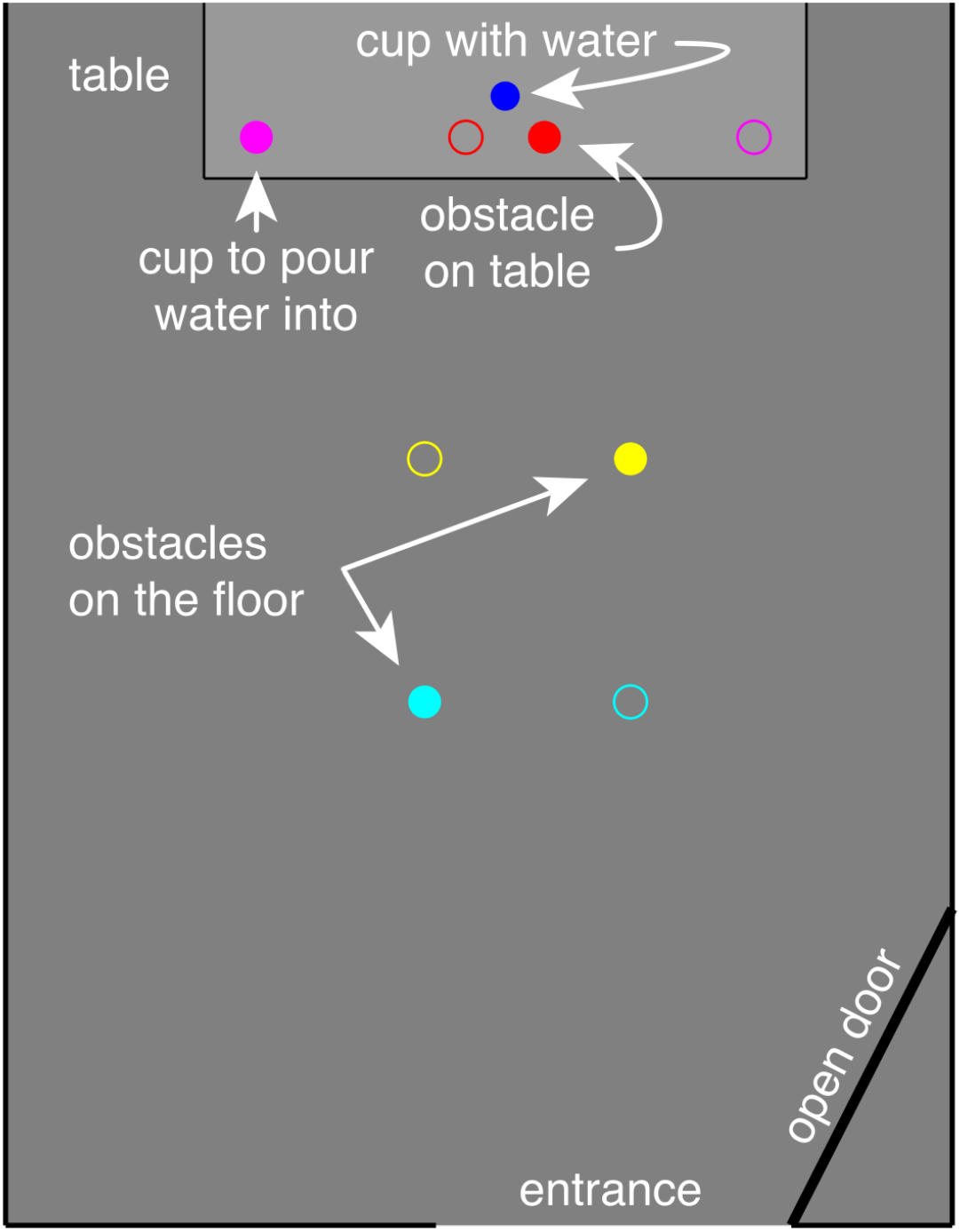
Schematic representation of a top view of the room, with the (open) door through which the participant entered the room at the bottom and the table at the top. The room extended beyond the top, but otherwise the representation is drawn to scale. The distance between the entrance and the near edge of the table was 280 cm. The task was to pick up the blue cup of water and pour its contents into the pink cup, while not knocking over the two obstacles on the floor along the path (turquoise and yellow cups), or the obstacle (red cup) near the cup of water. Open circles represent the alternative position of the obstacles and destination cups.

Participants wore a portable eye tracker (Pupil Invisible glasses, Pupil Labs, GmbH) that recorded their eye movements at 200 Hz and the scene at 30 Hz. The eye tracker consists of a spectacle frame holding the cameras that record the eye images connected with a cable to a smartphone, on which the data is stored. The scene camera is attached to the left side of the spectacle frame and has a field of view of 82 × 82 degrees. The eye tracker provides a good estimate of gaze position under variable conditions without requiring any calibration. A description of the glasses’ performance can be found at arxiv:2009.00508 or in Ghiani et al. (2023, 2024).

For each participant, we first started the recording of the eye tracker and then placed the recording phone in his or her pocket. Care was taken that the connecting cable would not hinder the participants’ walking and grasping movements. The participant started each trial by standing approximately 3 m from the door, facing away from the room. Meanwhile, one of the experimenters placed all the cups at the appropriate positions for the upcoming trial, and the other experimenter stood next to the participant holding a black card in front of the eye tracker’s scene camera. Occluding the scene camera made it easy to later segment the videos into separate trials (see *Data Analysis*). Once the configuration was ready, the experimenter removed the black card, which was the go-cue for the participant to start the trial. Participants then rotated their body and walked into the room to perform the task. No specific instructions were given other than that participants should grasp and manipulate the target cup with a right-hand precision grip and should be careful not to hit any obstacle and not to spill the water. Once participants had performed the task, they walked back out of the room to the experimenter with the black card and waited for the next trial to begin.

Each combination of the 16 possible configurations (2 near floor obstacle positions, 2 far floor obstacle positions, 2 table obstacle positions, 2 destination positions) was presented 5 times in a pseudorandom order for a total of 80 trials. Walking into the room and performing the grasping and manipulation task took approximately 12 seconds. Including walking back out of the room and waiting for the experimenter to set up the next configuration (and pour the water back into the target cup) a trial took about 22 seconds, so the whole experiment took about 35 minutes.

### Data analysis

The eye tracker provides information about gaze in the scene, but does not identify the items in the scene. Using differently coloured cups allowed us to identify the relevant items by their colour. Relying only on colour, light falling on the floor was sometimes mistaken for the yellow cup. To prevent this from happening, the software was designed to only look for cups at plausible positions relative to each other and to their positions on the previous image frame. Once the cups had been identified, the closest item to where participants were looking was determined for each frame of the scene image, as well as how far from that item gaze was directed. We also used the identified scene positions of the two floor obstacles, the target cup, and the cup into which the water was to be poured to estimate the position of the participant. This is not very precise, but it gives a reasonable idea of where the participant was.

We plotted the fraction of time that participants looked within 5 degrees of the center of each of the five items of interest (the two floor obstacles, the table obstacle, the target cup, and the destination cup). We also plotted the fraction of time that each of the five items was closest to the participants’ gaze, independently of how close to that item participants were actually looking. We estimated how far the participant was from the near edge of the table from the angular separation between the blue and purple cups. We combined this with the visible cups’ positions to estimate the participant’s lateral position in the room. We estimated where on the floor the participant was looking when the participant was looking at the floor as he or she entered the room by combining the known positions of the visible cups in the scene images with the participant’s eye height and estimated position in the room. For displaying the average paths that participants took to reach the table, we determined the median lateral position within 75 cm in depth for every 5 cm change in the distance from the table.

We also determined how long participants looked at the floor while walking toward the target object. We considered all times at which the participant was judged to be within 1 and 6 m from the table, and gaze was directed at the floor within the room and in front of the table. We determined this separately per trial and then averaged separately across the 16 possible configurations for each participant. To examine possible effects of the items’ configuration on the duration of gaze on the floor, we submitted those values to a 2 (near floor obstacle) x 2 (far floor obstacle) x 2 (target obstacle) x 2 (destination cup) repeated measures analysis of variance.

## Results

Our participants never knocked over any obstacle on the floor or table. Neither did they ever fail to pour the water into the destination cup without spilling any of the water.

We first examined which items participants were looking at as they approached the table. For this, we calculated the fraction of time during which their gaze was within 5 degrees of each of the five items of interest (Figure 2). At the beginning of the trial, before entering the room, participants primarily directed their gaze toward the center of the table, where the target cup and nearby obstacle were located. If both items were within 5 degrees of gaze, we assigned gaze to the closest item, but considering the resolution of the eye tracker and the proximity of these two items, it is not really possible to tell whether participants were looking at the target object or the nearby obstacle when participants were far from the table. To judge how much of the time gaze was directed at the combination of the target cup and nearby obstacle, one can simply judge the sum of the blue and red curves. Participants did also direct their gaze toward the destination cup and the obstacles on the floor, but they did so much less of the time, and mainly before entering the room. Thus, gaze was directed towards the regions that were relevant for planning how to pick up the target cup well in advance, rather than only being directed towards those regions after having made sure not to kick over the floor obstacles while walking towards the table. After entering the room, participants seldom looked at the floor obstacles, presumably because they had already established the positions of those items precisely enough to allow them to perform the task without having to look directly at them. As they approached the table, their gaze was mainly directed at the target cup and nearby obstacle. When the participant was near these two items, it became evident that they were mainly looking at the target cup.

**Figure 2.**
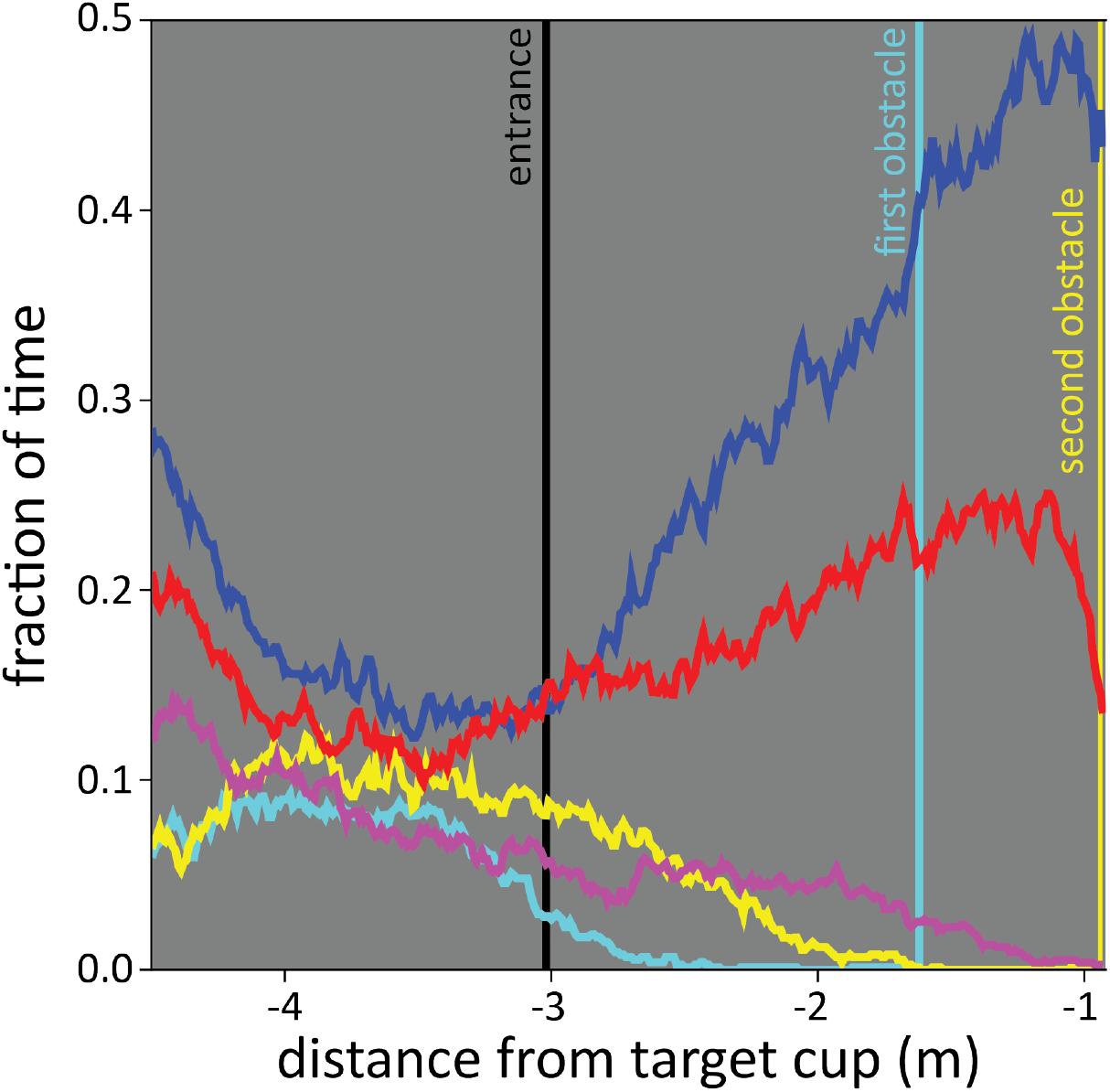
The fraction of the time that participants were looking at each cup as they approached the table. They were considered to be looking at a cup if their gaze was within 5 degrees of the centre of its image. Color-coding as in Figure 1.

Looking at Figure 2, it is evident that participants did not always look at one of the five items of interest (the sum of the five curves is lower than 1). Therefore, we also plot the fraction of time during which each of the considered items was the closest item to the participants’ gaze position. In that case, looking at the floor near the entrance is considered to be looking close to the first obstacle, and looking at the table surface is considered to be looking at one of the items on the table. Figure 3 demonstrates a similar pattern to the one depicted in Figure 1, but the peaks in gaze closest to the cups on the floor reveal that participants’ gaze shifted toward the floor when participants were about 2 m from the obstacle in question. As gaze was often closest to the floor obstacles, but not within 5 degrees of those items, we can infer that participants might have been looking where they plan to place their feet, rather than at the obstacles. Thus, they presumably rely on peripheral vision to ensure that they do not kick over the obstacles. In accordance with the impression given by Figure 2, once participants had entered the room, their gaze gradually shifted away from the floor toward the target cup and the obstacle next to it.

**Figure 3.**
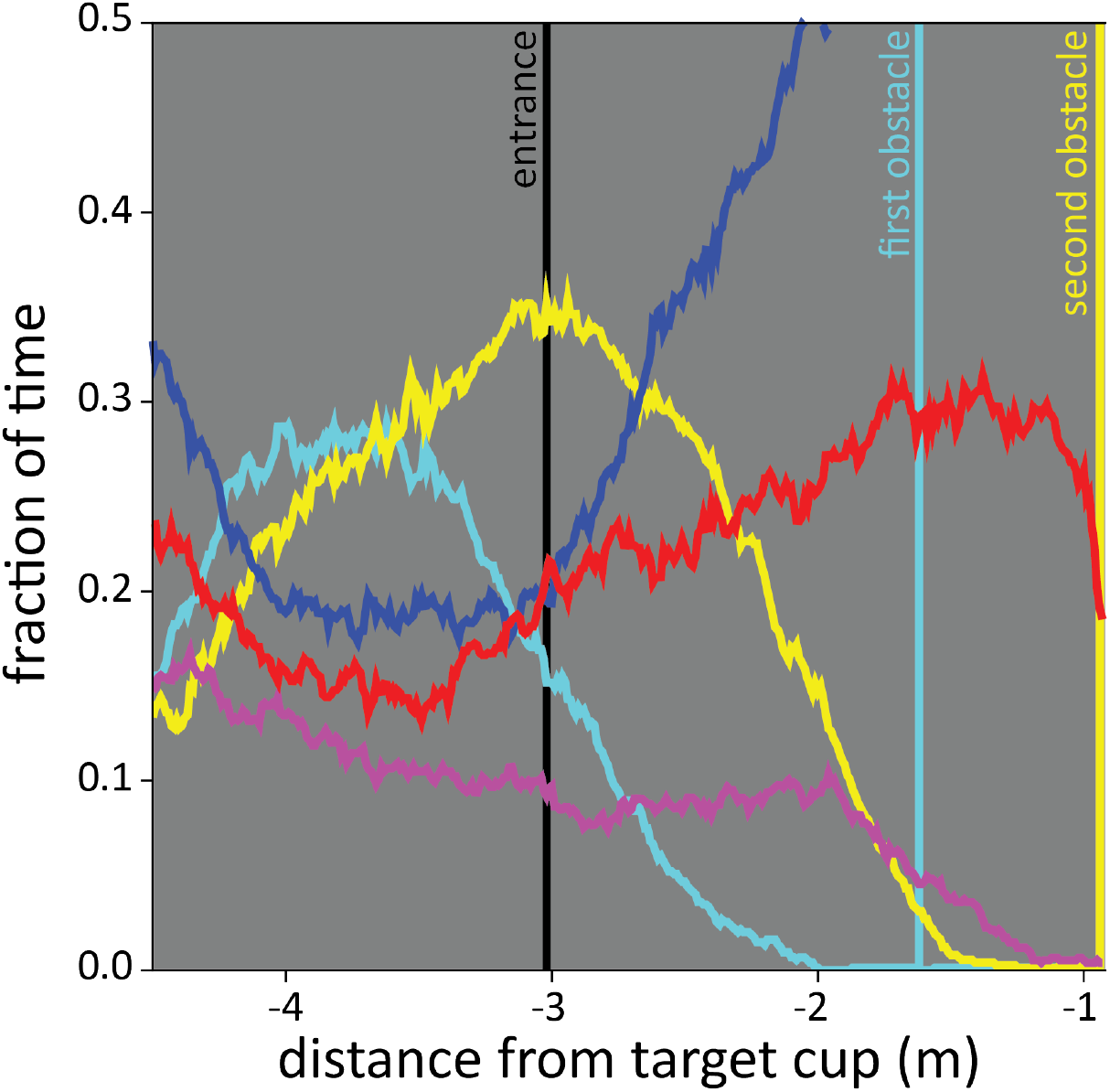
The closest cup to where participants were looking as they approached the table. Note that in this plot the five fractions sum to 1. Color-coding as in Figure 1.

That participants looked at their walking path rather than at the floor obstacles is confirmed in Figure 4. This figure shows the gaze positions on the floor together with the positions of the floor obstacles. The lines show the estimated walking path that the participants took (joining median lateral positions in binned distances from the table). All data was considered for this analysis as long as all the cups were at least partially visible (so they could be detected) within the scene camera’s images. This was usually so until the first (turquoise) obstacle on the floor disappeared below the scene camera’s view, but there are also some gaps in the data when the participant rotated his or her head too far to one side (so the purple cup was not visible). Moments at which gaze was not directed at the floor were ignored when plotting the gaze positions in Figure 4. Except when the obstacles were both on the left (leftmost panel), gaze was mainly directed at parts of the floor along the expected walking path (lines) that were close to the obstacles. Participants did not mainly look at the obstacles themselves. When both obstacles were on the left, participants’ gaze was less consistently directed towards a certain region of the floor. This might be because in this case it was relatively easy to walk to the table without knocking over the floor obstacles, because in this case the door and the target cup were both to the right of those obstacles so participants could just walk straight into the room. Overall, Figure 4 confirms what we inferred from comparing Figures 2 and 3: that when participants’ gaze was directed towards the floor, it was mainly toward critical positions for foot placement along their walking path, rather than toward the obstacles themselves.

**Figure 4.**
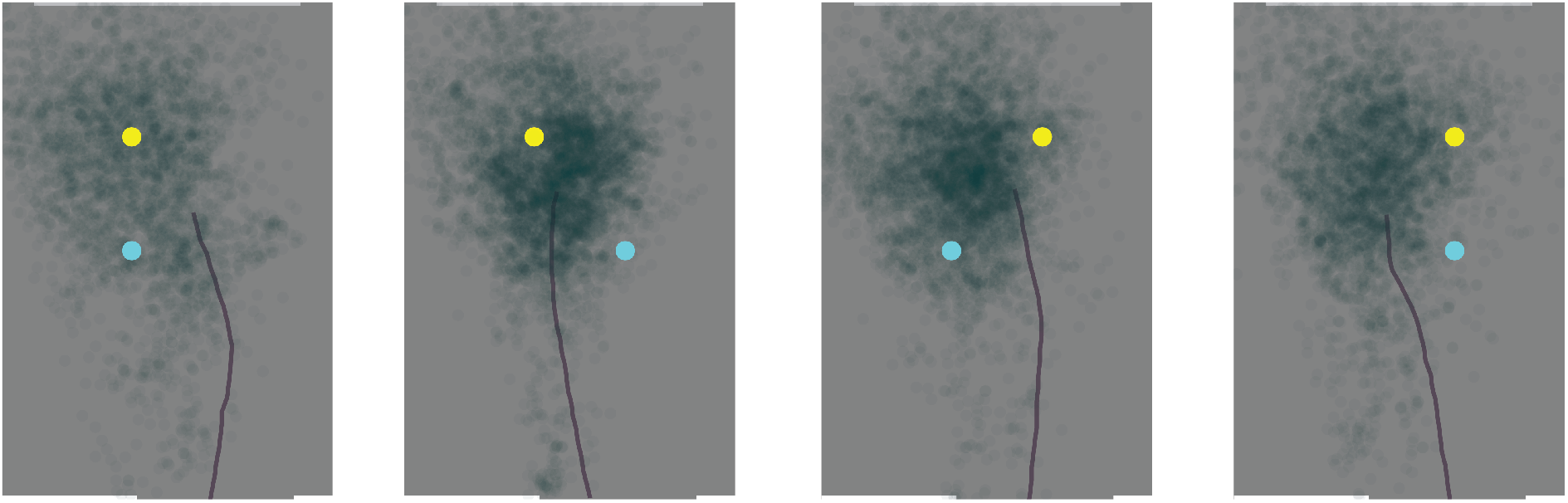
Estimated gaze positions on the floor (transparent green disks) and average walking trajectories (lines) for each of the four possible floor obstacle configurations. Plots show a top view of the room, from the door (bottom) to the table (top). The yellow and turquoise disks are the two obstacles.

If gaze is directed at the floor to guide foot placement at critical positions, one might expect people to look at the floor longer when the two floor obstacles were at opposite sides, because for those floor obstacle configurations participants usually walked in between the obstacles (see lines in the central panels of Figure 4). Moving one’s feet between the obstacles without kicking them over probably requires more precise guidance than just ensuring to move far enough to the side. This prediction is confirmed by a significant interaction between the two floor obstacle positions when comparing the time that participants spent looking at the floor (F_1, 9_ = 29.29, p = 0.0004; Figure 5). One might have also expected people to look longer at the target cup when the nearby obstacle constrained the grasping posture, and therefore to look shorter at the floor when the table obstacle was to the right than when it was to the left of the target cup. However, there was no main effect of the table obstacle position (F_1, 9_ = 0.076, p = 0.79).

**Figure 5.**
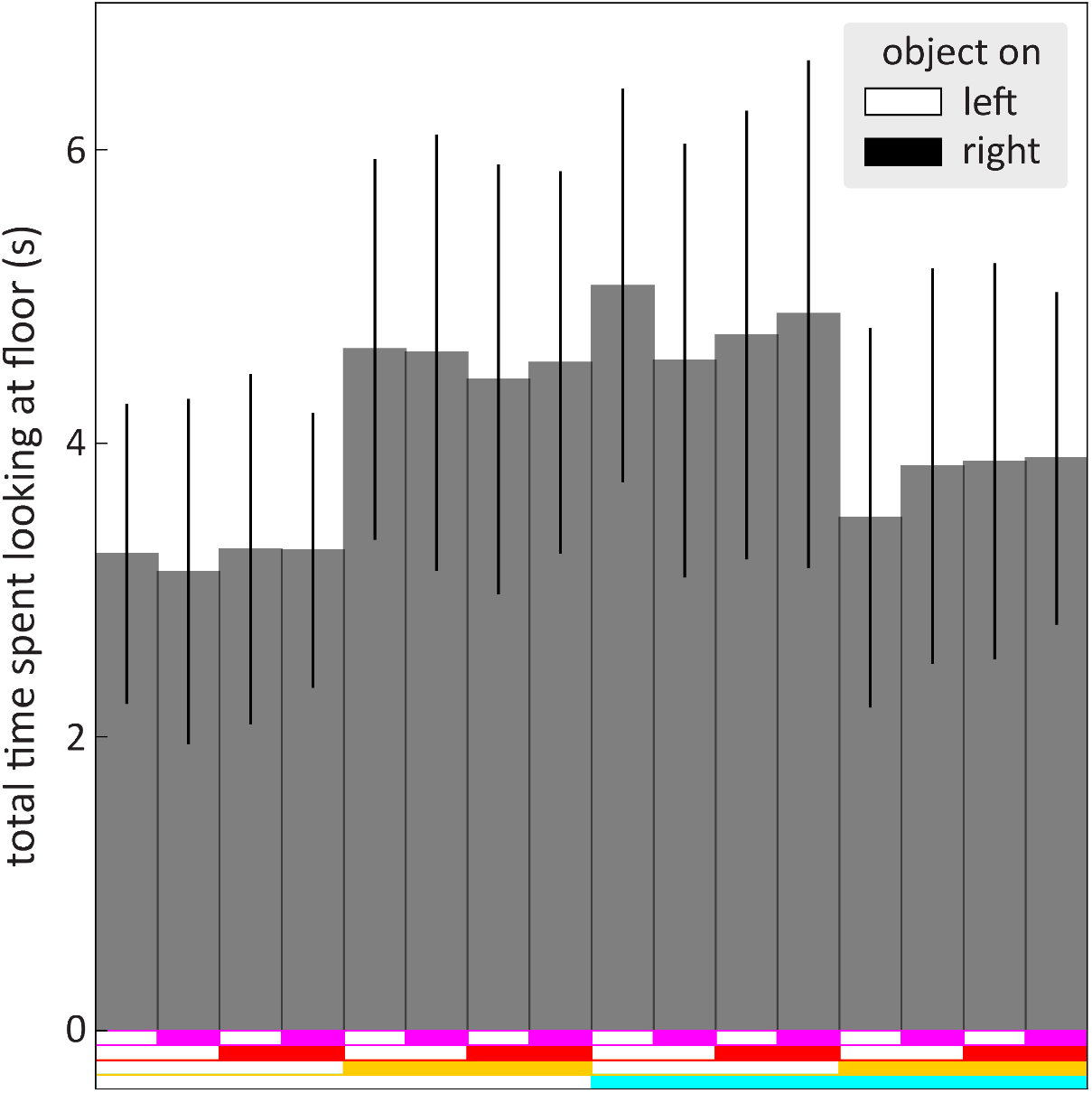
Total time spent looking at the floor for each of the 16 possible item configurations. Participants looked at the floor longer when the two floor obstacles (turquoise and yellow bars at the bottom of the figure) were at opposite sides (central 8 bars) than when they were at the same side (both on the left: leftmost 4 bars; or both on the right: rightmost 4 bars). Values are averages with 95% confidence intervals across participants.

In Figures 2-4 we averaged the data across all ten participants. However, there were clear differences between participants. This is shown in Figure 6, where each panel shows the closest cup to gaze (as in Figure 3) during all the individual trials of one of the ten participants. It appears that each participant has their own way of allocating gaze, which is rather consistent across trials. Some participants looked at the floor in front of them quite a lot as they entered the room (there is a lot of yellow and turquoise in the left half of the figure; e.g., bottom left panel). Others direct their gaze toward the items on the table from the beginning of the trial, with only occasional glances directed at the floor (e.g., top right panel). As the participants approached the table they mainly looked at the cup that they intended to pick up (a lot of blue at the very right of the panels of Figure 6). They also sometimes appear to have been looking at the obstacle near the target cup (shown in red), but since the two were very close to each other we cannot be very confident in allocating gaze between them. After the time period shown in Figure 6, when participants reached the table, they continued to look at the cup of water for up to one second while picking it up. They switched to looking at the destination cup while transporting the target cup toward it, and then looked almost exclusively at the destination cup, and possibly at the water being poured into that cup, for several seconds.

**Figure 6.**
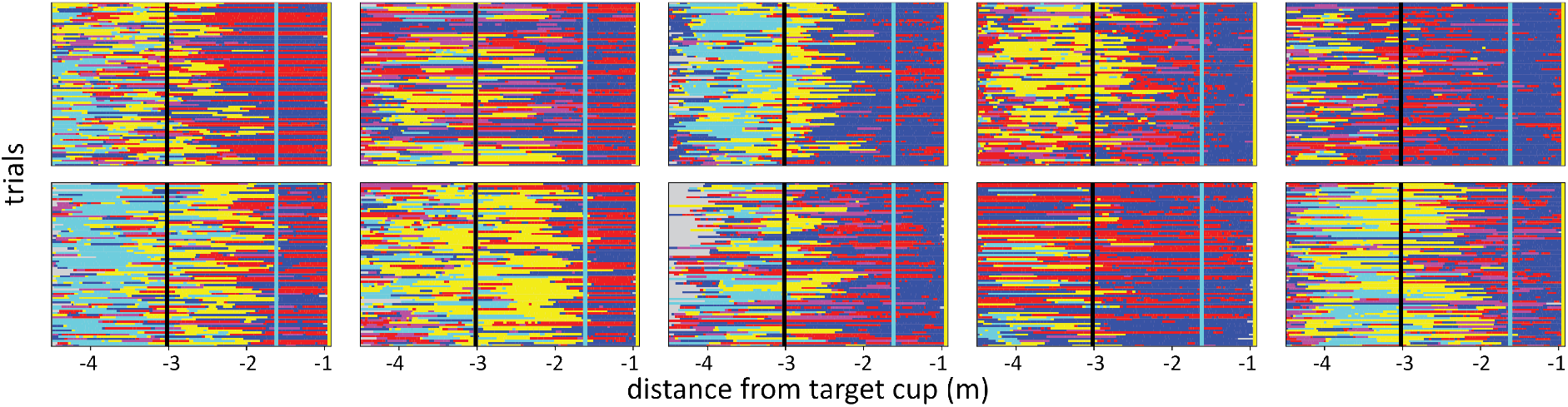
Closest cup to where participants were looking as they approached the table (as in Figure 3). Data for all trials. Each panel represents one participant. Each row represents one trial. Colors as in Figure 1. Grey sections represent moments in which we cannot judge where participants are looking because the cups are not within the scene image.

## Discussion

We were interested in how participants allocate their gaze when walking to grasp and manipulate an object if they have to avoid obstacles along their walking path as well as near the target object. We were particularly interested in whether participants would mainly look at the obstacles along their walking path as they enter the room, and only later look at the cup, as one might expect from some previous findings (Johansson et al., 2001; Patla & Vikers, 1997), or whether they would immediately mainly look at the object that is most important for the task (Rothkopf et al., 2007; Voudouris et al., 2016) or that requires the most precision: the cup of water that they have to pick up. Having an obstacle next to the cup of water made it more difficult to reach and grasp that cup, which might prompt participants to look at that cup and the nearby obstacle carefully to consider the best way to approach the table.

Participants looked at the most relevant positions in the room before entering the room. When looking at the floor, they mainly looked near where they would place their foot (Figure 4) several steps ahead along their walking path (Figure 3), despite there being nothing particularly informative to see at those positions, and those positions not being particularly salient. They seldom fixated the floor obstacles. It was already known that people look where they place their feet when this has to be done precisely, either because the task is to place them at certain places (Hollands & Marple-Horvat, 2001) or because the terrain requires precise foot placement (Matthis et al., 2018). In accordance with the latter finding, we found that participants looked at the floor more of the time when foot placement was more critical due to the configuration of the obstacles (Figure 5). But it is interesting to see that participants seldom fixated the obstacles when the only real constraint on foot placement was not to kick the obstacles. Instead they must have relied on remembered obstacle positions from brief glimpses earlier on, and on peripheral visual input as they approach the obstacles, to avoid kicking the floor obstacles over as they cross them while already looking at the target cup (Figures 3 and 6).

Long before entering the room, participants often already looked at the cups on the table (Figure 2). Knowing the cups’ positions might be important when entering the room, because determining the best place to stand for grasping the cup of water and even for later pouring its contents into the purple cup can be useful in determining how to approach the table. People are known to aim for certain hand and arm configurations at the moment of grasping (Voudouris et al., 2010; but see also Klein et al., 2021) and to consider obstacles in the paths of the fingers and of the arm when doing so (Voudouris et al., 2012). That is probably why people look at objects that they intend to grasp (Land et al., 1999; Voudouris et al., 2018) and at potential obstacles close to them (Grant, 2015) before initiating movements toward them. Here they do so well in advance.

When participants looked at the floor, they were most likely to direct their gaze toward positions near obstacles. They were most likely to look at such positions when they were about 2 m (about two steps) from them (Figure 3), which is consistent with earlier findings about looking where one will place one’s foot (Matthis et al., 2018). They looked where they intended to place their feet, as they did when faced with challenging terrain (Matthis et al., 2018), despite placing the foot not being particularly challenging in our study. In studies in which gaze was found to be specifically directed at obstacles, the obstacle was often on the path and had to be stepped over (Patla & Vikers, 1997). We therefore interpret our findings as support for the idea that gaze is primarily directed to the intended path, largely relying on peripheral vision for planning the path and avoiding obstacles. However, this need not always be the case. When people are specifically instructed to avoid obstacles, they do spend more time looking directly at obstacles (Rothkopf et al., 2007), especially if the position of the obstacle is uncertain (Tong et al., 2017), even if they can walk around the obstacles rather than having to step over them.

An obvious concern in any eye movement study, and one that is particularly relevant in studies performed under unconventional circumstances (while walking), is whether the measurements are accurate and precise enough to justify the analysis and conclusions. A very attractive feature of the Pupil Invisible eye tracker is that it provides quite precise measurements (Ghiani et al., 2024), even when walking (on stairs; Ghiani et al., 2023): a standard deviation of about 1 deg. However, the lateral placement of the scene camera gives rise to some systematic error. This was particularly relevant when looking at the cups on the table, because participants did so from distances ranging from about 6m to less than 1m. We corrected for this parallax error when evaluating gaze with respect to the cups on the table by assuming that gaze was estimated for a distance of 3 m. The reference distance of 3 m was found by assuming that participants did not look at the red cup (obstacle on the table) more frequently when it was on the left than when it was on the right of the target object. Finding the approximate distance at which gaze was no longer subject to such a parallax error will presumably have increased accuracy with respect to previous estimates (Ghiani et al., 2024). When looking at the floor, the viewing distance is often about 3 m (also considering the eye height) so such a correction is less important, and indeed when looking at steps systematic gaze errors were modest (not more than a few degrees; Ghiani et al., 2023). Importantly, most of the cups in our study were placed far enough apart to reliably be able to tell which of them participants were looking at. The only exception is that the blue and red cups were very close together, so until the blue cup had been picked up we cannot be very confident that we have always attributed gaze to the correct one of the two (as was already mentioned).

When relating gaze to the items in the image (the differently coloured cups) the main sources of uncertainty are the resolution of determining gaze and how precisely we can determine the cups’ positions in the images. The latter may sometimes be slightly wrong because, for instance, specular reflections are not considered to be part of the cup, but such errors will usually be very modest. Thus, the correct cup will probably usually have been identified when judging whether people are looking close to its centre (Figure 2), and the closest cup to where gaze is directed will generally have been judged correctly (Figures 3 and 6). More uncertainty is introduced when estimating the participant’s distance from the table from the distance between the blue and purple cups in the image, in combination with the actual distance between them in space of 66.5 cm. Errors in judging the cups’ positions in the image, small variations in where the cups were placed on the table (despite the clear markings), and small errors due to lateral variability in the participant’s position, will have given rise to some variability in the judged distance, which will have influenced the distances in Figures 2, 3, 4 and 6. Thus, some of the changes with distance might be slightly sharper than shown in those figures, but otherwise errors in judging the distance cannot account for any of the observed findings.

For Figure 4, we used the configuration of the cups in the image to estimate the participant’s position in the room, and used the judged positions of the participant and the cups, together with the participant’s eye height, to evaluate where exactly on the floor the participant was looking. Although we could not judge the participant’s position very precisely, the error in determining where they were looking on the floor is not too imprecise because the ordinal position relative to the cups is maintained. The only factor that is influenced by considering the participant’s viewpoint is the relative position with respect to the cups, which depends to some extent on where the participant is situated because it depends on the viewing angle with respect to the ground surface. Thus, we may be slightly overestimating the variability in gaze positions (Figure 4), but less variability would not change our conclusions. A final point that is worth mentioning is that we did not remove blinks (the data was collected before the Pupil Invisible provided automatic blink detection). Since the Pupil Invisible does give gaze estimates during blinks, and such estimates are included in our sample, they will have added (more or less random) noise to the gaze data. Thus, again, our analysis might be overestimating the variability in gaze. However, although our measures are certainly not perfect, we believe they are good enough to support our conclusions.

During the experiment, gaze shifted systematically between looking at different cups in a reasonable manner. The only exception is at the very beginning of the trials. When asked to prepare breakfast in an unfamiliar setting, people first look around to explore the available items and their positions (Hayhoe et al., 2003). Likewise, when entering a room for the first time, people first look around for relevant objects. They look around less, presumably using knowledge obtained the first time, when entering the same room again (Li et al., 2016). In our experiment, the positions of the obstacles and of the cup into which the water had to be poured changed between trials. Therefore, participants probably looked around to determine a suitable walking path before walking into the room.

When performing goal-directed arm movements, people usually look at the target object (Cámara et al., 2018; Danion & Flanagan, 2018; Grant, 2015; Voudouris et al., 2016), although they do look at obstacles when maneuvering a hand-held object around them (Johansson et al., 2001). When walking, people look at the ground more often when visual guidance of foot placement is more important (Hollands & Marple-Horvat, 2001; Matthis et al., 2018). We show that when walking into a room to grasp an object, people do not necessarily look at obstacles on their path to the object, but primarily look at the places at which interacting with the environment requires some precision, even if there is nothing in particular to see at those positions. Thus, people sequentially direct their gaze to where precision is required, relying on peripheral vision and memorized information to cope with challenges at other positions that are relevant for the task at hand.

## Acknowledgments

This study was funded by the Deutsche Forschungsgemeinschaft (DFG, German Research Foundation) – project number: VO 2542/1-1.

